# The Brain Image Library: A Community-Contributed Microscopy Resource for Neuroscientists

**DOI:** 10.1101/2023.12.22.573024

**Authors:** Mariah Kenney, Iaroslavna Vasylieva, Greg Hood, Ivan Cao-Berg, Luke Tuite, Rozita Laghaei, Megan C. Smith, Alan M. Watson, Alexander J. Ropelewski

## Abstract

Advancements in microscopy techniques and computing technologies have enabled researchers to digitally reconstruct brains at micron scale. As a result, community efforts like the BRAIN Initiative Cell Census Network (BICCN) have generated thousands of whole-brain imaging datasets to trace neuronal circuitry and comprehensively map cell types. This data holds valuable information that extends beyond initial analyses, opening avenues for variation studies and robust classification of cell types in specific brain regions. However, the size and heterogeneity of these imaging data have historically made storage, sharing, and analysis difficult for individual investigators and impractical on a broad community scale. Here, we introduce the Brain Image Library (BIL), a public resource serving the neuroscience community that provides a persistent centralized repository for brain microscopy data. BIL currently holds thousands of brain datasets and provides an integrated analysis ecosystem, allowing for exploration, visualization, and data access without the need to download, thus encouraging scientific discovery and data reuse.

## Introduction

In the era of big data and open science, the efficient management and sharing of research data have become crucial for scientific progress. Data availability promotes transparency and discovery by encouraging exploration and reuse. This has been a key goal of BRAIN initiative consortia including the Brain Initiative Cell Census Network (BICCN) which has collected thousands of brain microscopy datasets totaling several petabytes. BICCN aims to comprehensively catalog and understand the diversity of cell types in the brain including (i) understanding brain structure and function, (ii) cell type classification, (iii) mapping brain connectivity, (iv) creating a comprehensive reference cell type atlas, (v) advancing neuroscience research and (vi) data sharing and dissemination. BICCN generated numerous publications^1^, with several more expected over the next few years. Based on this initial success, the follow-on BRAIN Initiative Cell Atlas Network (BICAN) is shifting from predominately studying model organisms to focusing on primate brains. Consequently, the size of the data produced by BICAN is expected to eclipse that of BICCN as physically larger image volumes are acquired, including whole human brains.

The vast amounts of data produced by BICCN and BICAN efforts must be shared easily and promptly. To support the data-sharing goals, the National Institutes of Health (NIH) BRAIN Initiative^2^ established several archives to retrieve, store, and make available the data being produced by, or of interest to, the BRAIN Initiative (Table 1). Each archive houses modality-specific data, with the Brain Image Library (BIL) focused on optical microscopy. The mission of BIL is to be a public resource enabling researchers to deposit, analyze, mine, and share data by providing (i) a permanent repository for high-quality brain microscopy datasets, (ii) an analysis ecosystem with desktop visualization and high-performance computing (HPC) capability and (iii) user training, and support. BIL is housed at the Pittsburgh Supercomputing Center (PSC) and sits adjacent to the center’s flagship HPC system, Bridges-2^3^, a petascale resource that brings together HPC, Artificial Intelligence, and Big Data to empower scientific discovery. Proximity to HPC compute resources allows users to work with data using BIL infrastructure without the need to download terabytes of data, lowering the barriers and cost to the global scientific community. We believe BIL to be the largest archive of its kind and the first petascale brain microscopy data resource. Other related efforts include the Image Data Resource^4^, EBRAINS^5^, The Cell Vision^6^, The Cell Image Library^7^, and The BioImage Archive^8^. However, these resources are or were smaller in scale, limited in the size of the imagery they could accept, and did not provide the analysis ecosystem, thereby forcing users to download the data to be analyzed.

**Table 1.**
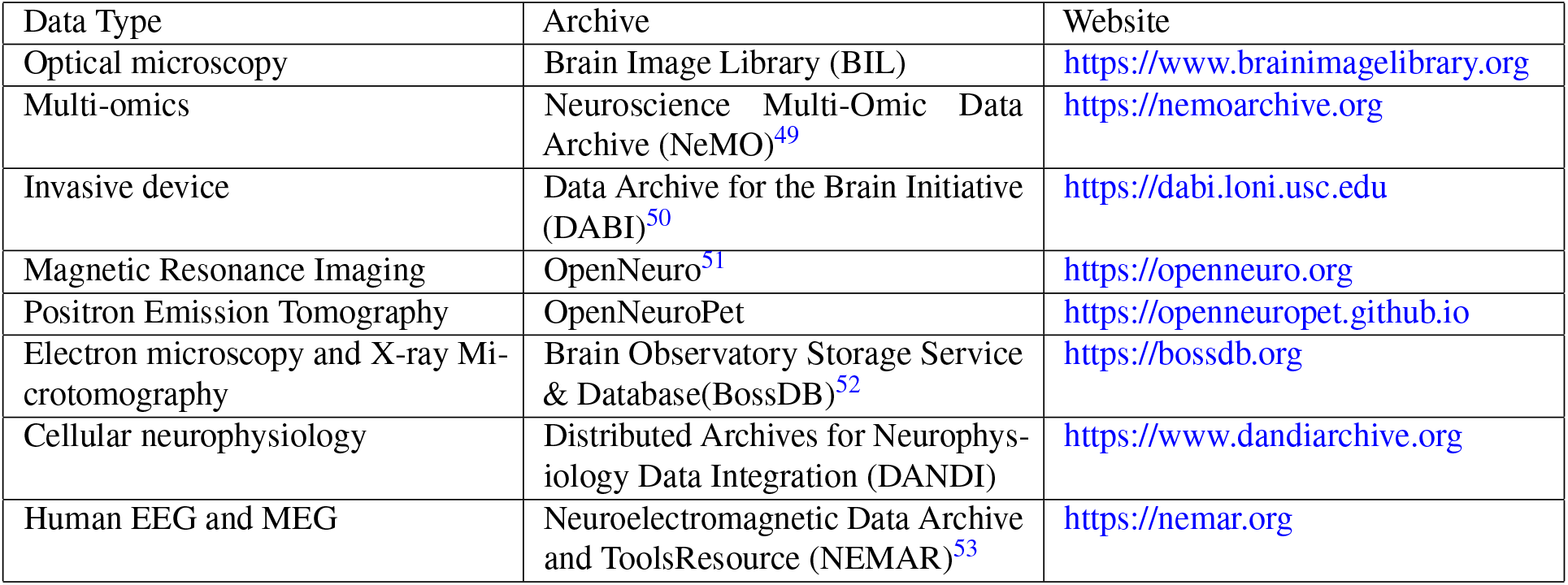
The BRAIN Initiative Data Archives and their data focus.

The scope of data accepted and available at BIL includes whole and partial brain microscopy image datasets, their accompanying derived data, and other historical collections of value to the community. BIL accepts optical image data that can include images directly from the imaging apparatus in a format that is open and accessible, and processed data that has been computed upon or transformed. Currently, BIL contains about 7,000 datasets representing over 268 data contributors. These contributors are from more than 45 unique affiliations and at least 6 different countries. This includes data from over 68 uniquely funded grants, 80% of which are BICCN. The datasets include high-resolution volumetric microscopy, cell morphology, connectivity, receptor mapping, cell counting/population, and spatial transcriptomics (Fig. 1a). Multiple species (mouse, marmoset, macaque, human, fruit fly, and ant) and a diverse range of microscopy technologies are represented including serial two-photon tomography (STPT)^9^, fluorescence micro-optical sectioning tomography (fMOST)^10^, light-sheet fluorescence microscopy (LSFM), and confocal microscopy, among others (Fig. 1b).

**Figure 1.**
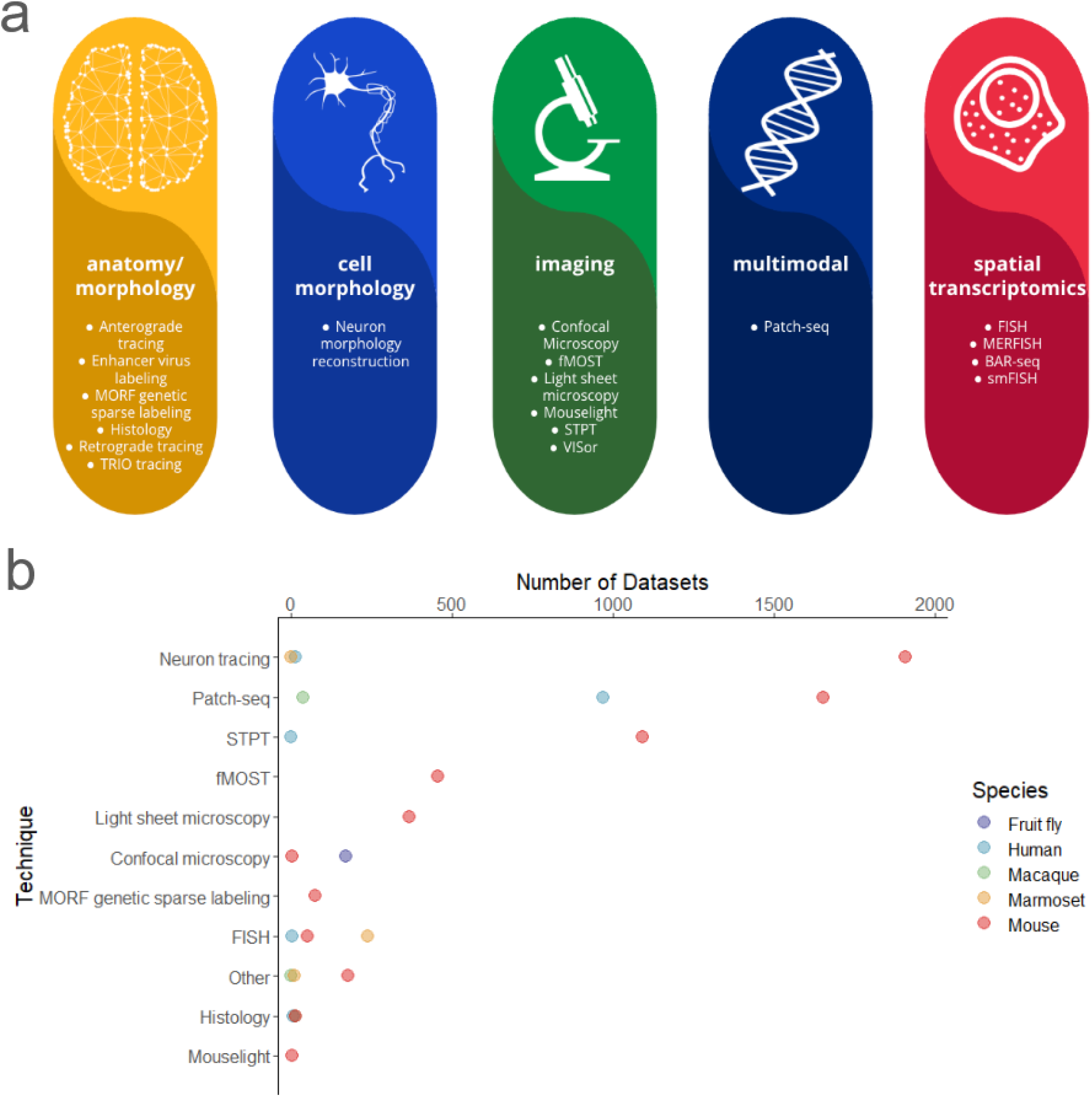
Summary of BIL datasets. a) An overview of the modalities and techniques of the data accepted and available at BIL adapted from the BICCN Data Catalog Glossary^48^. b) Distribution of datasets contributed to BIL by technique and species at the time of writing. The code and data used to generate panel b can be found at doi:10.6084/m9.figshare.25213781. The same code can be used to generate a panel for the most recent state of BIL datasets by downloading the summary metadata linked at the BIL search portal or directly [here](https://submit.brainimagelibrary.org/search/summarymetadata). Each data point is representative of an n=1.

## Methods

BIL provides researchers with a platform to share their data, access and analyze public data, collaborate with other researchers, and publish their findings. Data contributed to BIL is distributed in a way that encourages the broadest use and reuse. Data storage and computation are free. There is no limit on the data size, number of datasets uploaded per investigator, or the duration of data storage. The BIL team designs and maintains the infrastructure underlying the resource which allows BIL to respond flexibly to the needs of the neuroscience community.

### Image file formats

Historically, BIL has accepted data formats that are broadly reusable by the community. Most deposited images are whole brain volumetric stacks in native TIFF and JPEG 2000 image file formats. Currently, we are encouraging the use of high-performance next-generation file formats (NGFF)^11^ such as multiscale OME-Zarr^12^, which include multi-resolution image pyramids optimized for rapid visualization and scalable analysis. While BIL continues to accommodate historically accepted file formats, datasets in these formats may not leverage the complete range of visualization capabilities now provided by BIL as discussed below.

### Metadata

To maximize the value and impact of data, it is essential to ensure that datasets are Findable, Accessible, Interoperable, and Reusable (FAIR)^13^. The implementation of FAIR standards requires standardized metadata, which offers a detailed structured description of the data. Detailed standardized metadata promotes data reuse and facilitates the reproducibility of scientific experiments. It is especially important for peta-scale data repositories, where data users must be able to quickly understand the context and limitations of the datasets before investing time and resources needed to examine and analyze them.

BIL initially started with a simple metadata model that included 14 flexible fields for broad descriptions of datasets. In 2021, the initial model was upgraded to the 3D Microscopy Metadata Standards (3D-MMS) through a BRAIN Initiative standardization project^14^. This standard is in the process of being adopted by the International Neuroinformatics Coordinating Facility (INCF)^15^. The 3D-MMS model is aligned with DataCite metadata^16^ and includes information about the data contributors, project funders, and associated publications. In addition, the model contains descriptive metadata about the dataset, the specimen, and the instrumentation that was used to generate it. There are also 41 fields within the standard that align with the OME metadata schema^17^. Recently the 3D-MMS were expanded to include new fields for better descriptions of neuron tracing datasets deposited at BIL.

### Finding and downloading data

At BIL, data findability is ensured by assigning several unique identifiers. Principally, a DOI is issued to each dataset upon publication. This provides a unique stable identifier for easy tracking of data use and simple citation. Each dataset has a landing page (example doi:10.35077/ace-cab-ear) that provides detailed metadata, data descriptions, related datasets, and publications. All of the persistent identifiers issued by BIL include:

- <u>Submission Identifiers.</u> A unique ID is assigned to all data that is part of a submission to the archive. The submission identifier is a 16-digit hexadecimal string. An example of a BIL submission identifier is abcdef0123456789.
- <u>Dataset Identifier.</u> All datasets are assigned unique identifiers. These identifiers consist of a string of (English) pronounceable triplets. An example of a BIL dataset identifier is “ace-cot-new”.
- <u>Traced Neuron Identifier.</u> BIL assigns unique identifiers to traced neurons when additional metadata is available for the neuron. These identifiers are similar to Dataset Identifiers with the exception that they all begin with the triplet “swc”.
- <u>DOI (Dataset).</u> When a DOI is issued for a BIL dataset, it will be assigned a URL with the BIL doi prefix (10.35077) plus the data identifier. An example of a BIL dataset DOI is: https://doi.org/10.35077/ace-and
- <u>DOI (Collection/Group).</u> Upon request, a DOI can be issued which groups datasets together into a collection. For example, all datasets associated with a publication can be grouped into a single DOI. These DOIs begin with a “g.” followed by a number. An example of a BIL group DOI is: https://doi.brainimagelibrary.org/doi/10.35077/g.948

#### Inventory search

A metadata API and web portal that uses the API are available to search for datasets and return metadata along with links to the data. The metadata API is capable of full-text searching, including all individual metadata fields. The API has two primary user-facing endpoints:

- *query* - the query endpoint will query the system and return a JavaScript Object Notation (JSON) document with all matching dataset entries.
- *retrieve* - the retrieve endpoint will return the metadata as a JSON document for the given Dataset Identifiers.

In addition to the API, BIL provides a web search interface that can query the metadata API in a variety of different ways, and display selectable results based on the query.

#### Bulk download

While individual links to data are available through the web search interface described above, all BIL data is available for authenticated access using standard Unix tools (rsync, sftp, scp, etc.). Access without authentication is available through web URLs (https://download.brainimagelibrary.org) and a Globus/GridFTP endpoint “Brain Image Library Download”. To facilitate the download of individual datasets, each dataset has an associated manifest file in JSON format which contains checksums and related metadata. Downloading data from BIL is optional as BIL provides visualization tools and a comprehensive analysis ecosystem (described below), which includes a wide range of computational resources and software to explore data in place.

### Data submission process

Data generators interested in submitting their own data to BIL can do so by following the data submission process outlined in Fig. 2a. The submission process includes (i) data upload through the [submission portal](https://submit.brainimagelibrary.org/), (ii) file validation, (iii) curation of data and metadata by a dedicated data curator, and ultimately, (iv) the publication of the data. Data submitters have the option for a limited embargo period for their data. During the embargo period, only the associated metadata will be accessible to the public. BIL provides a submission portal with an administrative view for project management. The portal offers a comprehensive overview of the data submitted by multiple contributors within the research project or subproject along with the submission status, and provides a bridge to link metadata with uploaded data. Data can be deposited over the network, using Linear Tape-Open tapes, or with portable shippable drives. The majority of data contributors transfer data over the network. BIL’s unique network analysis and diagnostic support helps to resolve transfer bottlenecks between the BIL data center and the contributing site.

**Figure 2.**
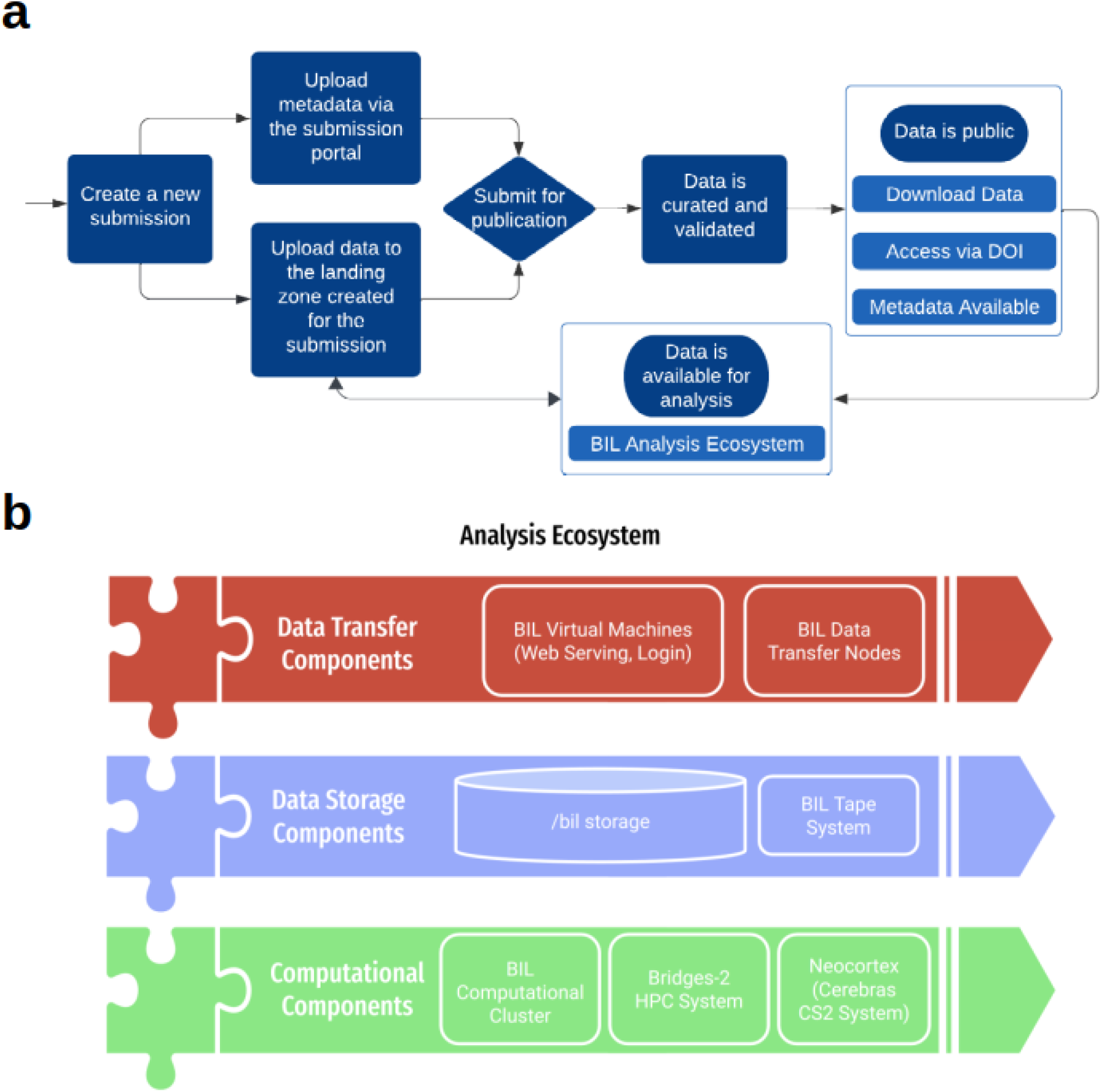
BIL workflow and resources. a) The BIL resources are utilized in the comprehensive data submission and publication process, as depicted in the flow diagram. b) This process involves data transfer (top), storage (middle), and computation (bottom) within the BIL analysis ecosystem.

### BIL analysis ecosystem

#### Architecture

The BIL Analysis Ecosystem offers computational resources designed for visualizing and processing BIL data. This ecosystem is equipped for desktop visualization and high-performance computing to handle pre-submission data processing and post-submission exploration. The architecture seamlessly integrates mass storage, networking, and HPC components within a unified environment (Fig. 2b). Data transfer components allow the capability to efficiently support large file and data transfers. The data transfer infrastructure includes redundant data transfer nodes integrated with wide area and local area network connections. The dedicated system provides a remote desktop environment to run interactive applications for visualization and analysis such as Fiji^18^, Napari^19^, and Vaa3d^20^, and to serve web portals dedicated to tasks such as data ingestion, and APIs. The data storage components consist of a fault-tolerant Lustre multi-petabyte scalable filesystem which is accessible on all of our computing platforms. Tape is used for making archival backups. Computational components include a variety of large memory nodes, GPU nodes, and access to high-performance computing resources, including PSC’s Bridges2^3^ and Neocortex^21^ systems for extensive data exploration. More thorough technical architecture description of the ecosystem is given in a dedicated publication^22^.

#### Software

Various software options are available for users, including traditional software development languages, higher-level scripting languages, and applications. Most application software at BIL uses Lmod modules^23^ that provide a uniform and stable method to access multiple software versions for reproducible workflows. Software available through the module system at BIL include image analysis tools, machine learning toolkits, file conversion tools, and other open-source packages. BIL staff are available to work with users to enable new open-source software upon request.

#### Resource management

The ecosystem uses the SLURM (Simple Linux Utility for Resource Management^24^) scheduler. SLURM is widely used in the field of scientific computing in HPC clusters due to its flexibility and scalability. It provides a robust framework for job submission, scheduling, and resource management, enabling multiple users to effectively share cluster resources. At BIL, SLURM can be used interactively and for batch processing. Access to resources is provided through login nodes which are capable of allocating resources on large memory nodes with up to 4 terabytes of random access memory, GPU nodes with up to 8 GPUs, and thousands of smaller memory nodes.

#### Containers and workflows

Containerized software organized into workflows is the best way to ensure reproducibility of analysis and interoperability of tools in different computational environments. Singularity^25^ containers are supported on the analysis ecosystem. Singularity is a standard tool on HPC clusters that provides a secure way to create reproducible applications. While users can create their own containers, BIL provides several public singularity definition files, which users will be able to use as models for their own containers or pull for use on local infrastructure or public cloud computing environments. Most existing Docker containers can be converted to Singularity containers. In addition to containers, workflows developed using the Common Workflow Language (CWL)^26^ standard or SnakeMake^27^ can be run on BIL systems.

#### Web computational gateway through Open OnDemand

The ecosystem also supports an OpenOnDemand (OOD)^28^ [instance](https://ondemand.bil.psc.edu/) to explore and compute on data without leaving a web browser. OOD is an intuitive, innovative, and interactive web-based interface to remote computing resources. OOD enables users to connect to BIL resources through in-browser remote desktop, write code using Visual Studio Code, create and run python scripts through Jupyter^29^ and R scripts through RStudio.

### User support and training

BIL provides office hours regularly (the schedule is posted on BIL [website](https://www.brainimagelibrary.org/contact.html)). For general inquiries, user support is also offered through an email helpdesk at. Typical questions received are about data usage, tools, or software. Data submission workshops are half-day hands-on online sessions that are provided regularly for new data contributors. These workshops outline all aspects of the data publication process including data submission, data structure, and metadata submission and offer practical assistance to users. The Data Ecosystem workshops involve tutorials for running analysis jobs at BIL and using tools such as SLURM, Open OnDemand, and Jupyter Notebooks to interact with datasets and images housed in the archive. All workshop materials are available at BIL GitHub. The recordings are available on [YouTube](https://www.youtube.com/channel/UCgUUJXxaeuEmozzUIaGmTuQ).

## Results

The Brain Image Library is a hub for neuroscientists to store, visualize and analyze data. They can explore thousands of public datasets and collaborate in a uniquely resource rich environment. BIL is designed to be equally useful for novice users and experienced computer scientists. Users have access to BIL’s fully featured analysis ecosystem, which offers everything from standard command line access to remote desktop environments. In a continued effort to make data more accessible to all users, BIL has developed tools for easy visualization of terascale datasets over the internet without the need to register for an account or be an HPC expert. In addition, we have curated examples using graphical online tools like Jupyter notebooks that scientists may adapt for their own usage. Below we explore and share examples of these visualization and analysis tools.

### Visualization

The microscopy data stored at BIL is inherently visual. Visualization is crucial to the scientific process, as it provides rapid validation of experiments, presents feedback on the appropriateness of feature extraction algorithms, facilitates understanding of the data content and ultimately encourages post hoc reuse by the scientific community. Additionally, visualization is an excellent educational tool for teachers and students who wish to explore brain structure, compare and contrast imaging modalities, and understand the scientific process. The inherent beauty of microscopic datasets should not be overlooked as a motivating factor to propel students into science, technology, engineering, arts, and mathematics fields and compel citizen scientists to get involved in the process of discovery. However, the size and diversity of imaging data at BIL can pose a stumbling block for visualization. To address these concerns, we use Napari and Neuroglancer to visualize data over the internet without the need to download. In the case of Neuroglancer, datasets can be visualized in a common web browser, even using a smartphone.

#### Napari

Napari^19^ is an open-source multi-dimensional image viewer integrated into python ecosystem, that can efficiently load large image stacks or multi-scale data. We developed a plugin for napari called napari-bil-data-viewer^30^ that enables users to visualize publicly available datasets at BIL over the web without registering for a BIL account, thereby making the data more easily accessible and reusable in line with FAIR^13^ principles. The napari-bil-data-viewer can be installed on a low-end laptop, and the software requirements and technical expertise required are minimal. The current version of the plugin is pre-loaded with a comprehensive set of downsampled (summary) whole brain fMOST datasets (Fig 3, left) and neuron morphology SWC^31^ files associated with these brains. Additionally, arbitrary image stacks, multi-scale data in OME-Zarr format, and neuron morphology files can be visualized by providing a URL. Napari goes beyond simple visualization. Since the data are loaded as standard napari layers, they can be used as input to other plugins from the rich napari ecosystem, which includes tools for imaging, image analysis, and which cater to the needs of neuroscientists.

**Figure 3.**
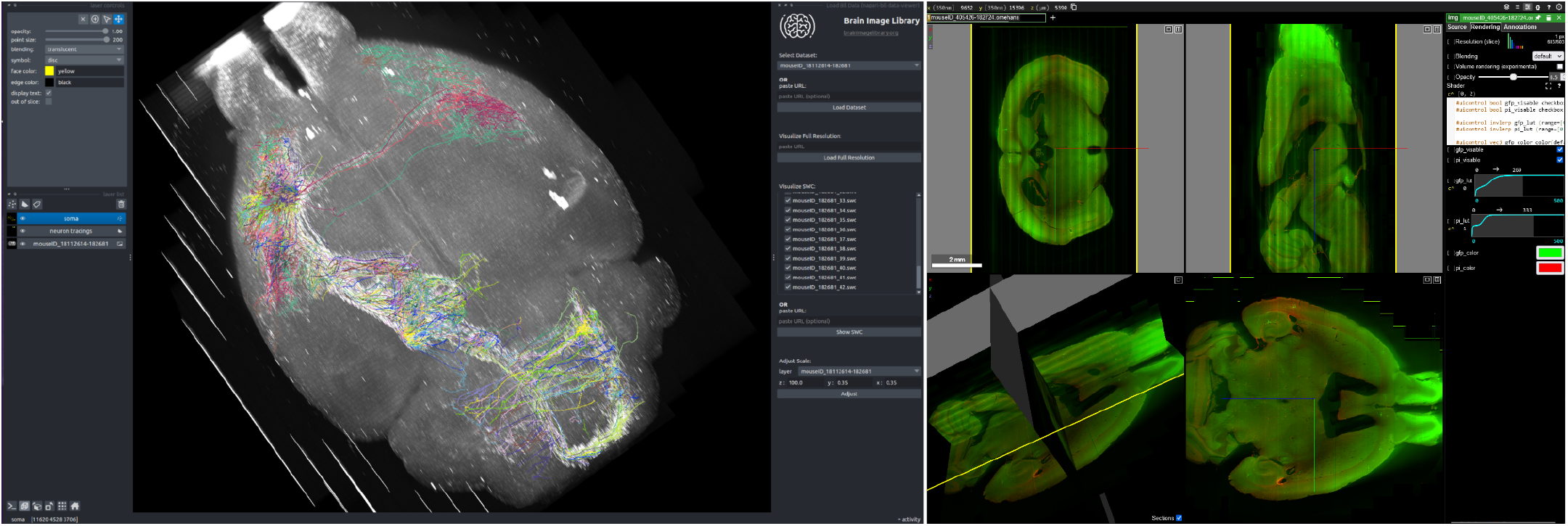
Visualization of BIL data. The bil-data-viewer plugin for napari (left) displays a 3D view of a fMOST dataset with overlaid neuron morphologies rendered from the SWC files which are shown along with the plugin interface. Neuroglancer visualizations (right) are available for many BIL datasests. A link to the neuroglancer example shown here is available at the following [search url](https://api.brainimagelibrary.org/web/view?bildid=ace-bag-kit) and is generated from the dataset located at doi:10.35077/ace-bag-kit. The neuron morphology dataset shown in this example is available at doi:10.35077/web, and the fMOST images are available at doi:10.35077/ace-ban-ear. These images were generated to illustrate the capabilities of the napari-bil-data-viewer and BIL neuroglancer viewer. The image on the left can be reproduced by installing napari-bil-data-viewer (v0.5.1 and napari v0.4.18). Image acquisition tools and parameters can be found in corresponding metadata at BIL. The lookup tables were adjusted minimally, for the entire image.

#### Neuroglancer

Neuroglancer^32^ is a popular open-source 3-dimensional image viewer created and maintained by Google and the broader community. Neuroglancer is widely used by the neuroscience community, including the customizations^33,34^ made by different labs meeting their specific needs. These needs include the ability to quickly and easily visualize cloud-based datasets, overlay annotations, and share specific views of the data (camera parameters, lookup table settings, zoom, etc). Neuroglancer scales well for visualizing petascale datasets. Several multiscale file formats are supported including the latest OME-Zarr standards. As discussed below, BIL can represent data from other multiscale formats (for example, Imaris^35^) as OME-Zarr or neuroglancer precomputed on the fly, without duplication of data. This allows BIL to make neuroglancer compatible with multiple image formats that would otherwise not be supported. Users can visualize arbitrarily large datasets even on a smartphone, at the speed of mobile internet, without registering for an account at BIL or downloading data. Fig. 3 (right) shows an example of visualizing BIL data with Neuroglancer.

### Data analysis

BIL serves a diverse community of users, using a variety of model organisms, imaging modalities and experimental techniques. We support individual groups to understand their analysis needs, guide them towards appropriate software and implement new tools as requested. Neuroscience is a diverse scientific discipline and there is a large and rapidly growing list of community derived tools. As such there is no universally agreed upon standard or workflow for analysing light microscopy brain images, especially for the recent advancements in whole brain images^36^. However, for common tasks such as registration to the brain atlases and mapping labeled cells, the Brainglobe family of tools^37^ provides an excellent starting point. The Brainglobe atlas API^38^ provides a universal way of accessing a number of atlases for different model organisms and imaging modalities. Registering raw brain images to an atlas is accomplished through Brainglobe brainreg^39^. Atlas registration ultimately allows results from different brains to be combined and compared in common atlas coordinates.

Here we provide an example pipeline based around the Brainglobe tools which is applicable to our largest corpus of data, STPT and fMOST image stacks. Figure 4 shows detection of labeled soma in a whole fMOST (panel a) and STPT (panel b) brain, and registration of these brains to the Allen mouse brain atlas^40^ (panel c). As a benchmark, registration of a whole STPT mouse brain at resolution (z, y, x) = (50, 1, 1) microns, to the 10-micron Allen atlas using one of the BIL large-memory nodes took about 90 minutes, whereas downsampling and registration of a 6-Terabyte full-resolution (1, 0.35, 0.35 microns) fMOST brain took around 6 hours. The processing times will vary depending on resources requested and availability of resources (i.e. resources are not in use by other users) at that moment. In panel d, comparison of cell distributions from two different STPT brains are displayed in atlas space on an interactive 3D model provided by Brainglobe brainrender^41^. The pipeline used to produce these results is publicly accessible and embedded in an interactive jupyter [notebook](https://github.com/brain-image-library/workshops/blob/4e11e8d18cd5df9d3e13b33a12e14a6d06ad7546/bil_pipelines/brainmapper.ipynb) which can be run from the BIL OpenOnDemand web [portal](https://ondemand.bil.psc.edu/) after registering for a BIL account. We demonstrated the use of these tools at our interactive workshops, for which the recordings are available on the BIL [YouTube channel](https://www.youtube.com/channel/UCgUUJXxaeuEmozzUIaGmTuQ), and the source code is available on github. Other examples of workflows have been demonstrated at the brain registration [hackathon](https://github.com/InsightSoftwareConsortium/GetYourBrainTogether/tree/main/HCK01_2022_Virtual), which brought together experts in the field in an effort to create reproducible image registration workflows.

**Figure 4.**
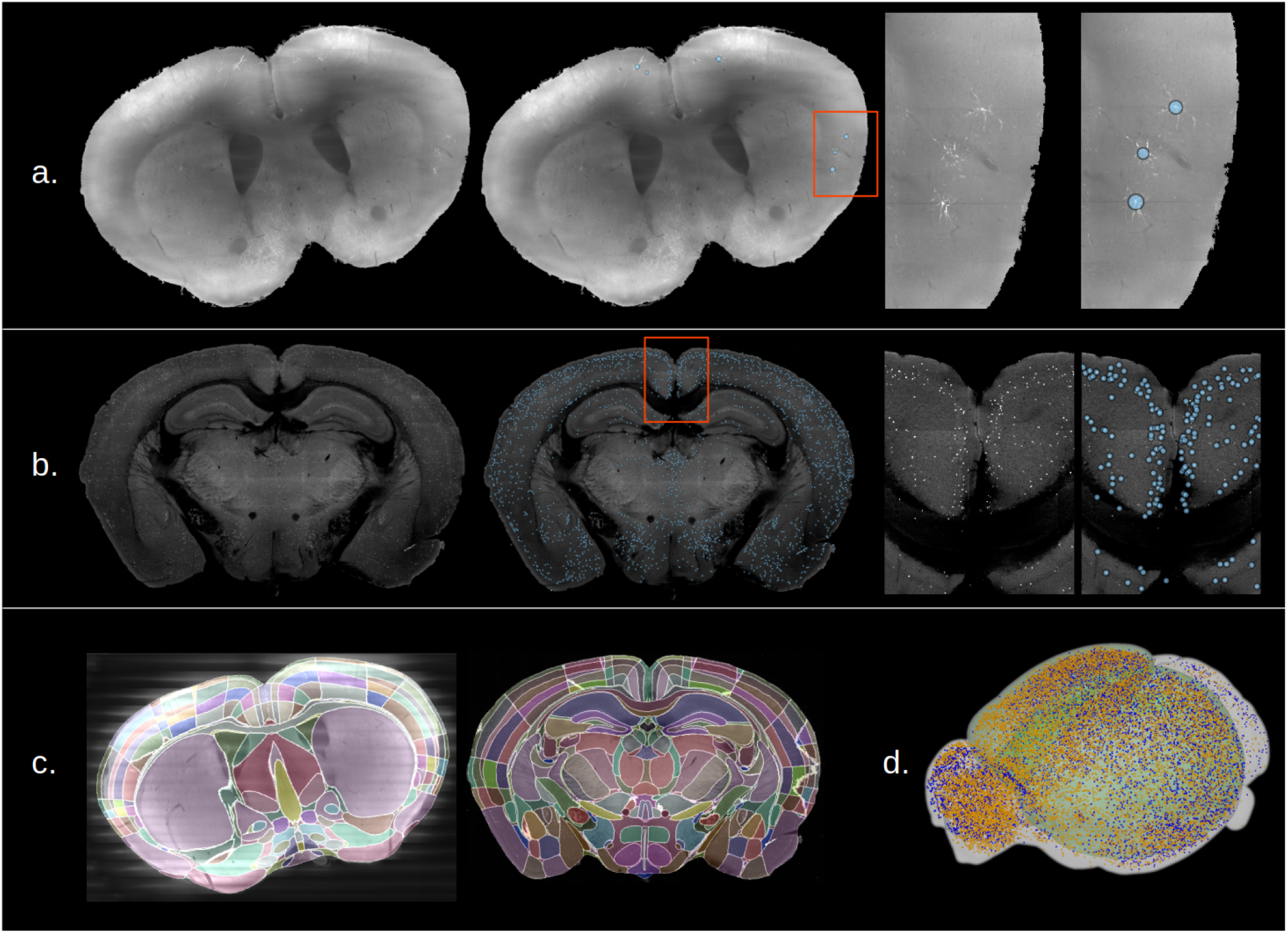
Analysis of whole brains deposited at BIL using [Brainglobe](https://brainglobe.info/) pipelines. a) Detection of viral labeled neurons in a whole fMOST mouse brain (BIL ID: ace-add-lit). b) Detection of labeled neuronal somata in a whole STPT mouse brain (BIL ID: ace-age-cop). c) Result of registering the two above brain images to the Allen mouse brain atlas. d) Comparison of the locations of tdTomato expressing neurons in P56 mouse brains from a vasoactive intestinal peptide cre-driver line. Displayed are points from male (yellow, BIL ID: ace-dub-how) and female (blue, BIL ID: ace-dub-hot) mice displayed in Allen atlas coordinates. Every tenth cell is shown for image clarity.

## Discussion

Here, we presented the Brain Image Library, a unique microscopy data resource that equips the neuroscience community with an extensive collection of brain images representing a variety of microscopy-enabled experiments. Uniquely, BIL provides a platform for analyzing and sharing large-scale imaging data that would otherwise be difficult to access. BIL supports data generators by preserving valuable experimental data and offering them free resources for analysis. To facilitate data reuse, the visualization and analysis ecosystem enables data to be viewed over the web and analyzed in place on HPC systems. These essential features become even more vital as the collection of microscope-enabled experiments of human brain data is deposited by efforts like BICAN. BIL is well-positioned to accept larger and more complex data as the neuroscience community continues to innovate. Indeed, we encourage all members of the neuroscience community to deposit their microscopy data in BIL. Below, we discuss pressing needs and directions of our ongoing and future work in serving the neuroscience community.

### Image format diversity

Collaborative experiments can produce a multitude of formats and data structures that ultimately present challenges for data interoperability and reuse. There are hundreds of imaging data formats currently in use, including standardized formats alongside vendor-specific formats. To integrate data effectively, researchers must possess domain-specific knowledge of the imaging formats in which the data is stored. Moreover, analysis tools often support a limited number of file formats^36^, which will require that the data be available in the formats selected by the tool developer.

To address this, community-driven efforts like the open microscopy environment (OME)^17^ have developed excellent tools like Bio Formats^42^ with support for about 150 file formats. However, direct integration of Bio Formats into image processing applications is relatively rare, and it is most often used to convert data from non-standard to standard formats post hoc, essentially duplicating it. BIL provides access to Bio Formats if traditional conversion is required. However, as hundreds of terabytes to petabyte-sized primate brain images start to be deposited in BIL, this approach becomes extremely costly and impractical in terms of the computational and storage resources required.

To address data format challenges and avoid data duplication, we have been developing an on-demand transformer. This transformer facilitates the efficient delivery of data residing on BIL file servers to end-users. Data can be delivered for local computing or over the internet in various formats. Thus, we aim to provide flexibility by delivering data in useful ways while maintaining storage efficiency within BIL. The objective use of the one-demand transformer is twofold: visualization and data interaction. Arbitrary requests at the voxel level can be made, ranging from individual voxels to entire datasets. Since data can be delivered in standardized formats, it is compatible with visualization tools like Napari or Neuroglancer and can also be used by programmatic interfaces for data analysis. For instance, it allows the extraction of specific pixel coordinates within multiple resolution versions of a multi-terabyte whole brain dataset, allowing for the extraction of regions of interest at different scales. Although still in development, this capability is currently being used to serve our Neuroglancer data viewer described in the results section. The application currently offers single-click access to over 1 petabyte of public volumetric imaging data from over 80,000 files. Data can currently be served as multiscale OME-Zarr and neuroglancer pre-computed formats for visualization or for computing in any application compatible with these formats. Currently, we are in the process of converting legacy datasets into multi-scale OME-Zarr. However, we are encouraging data providers to submit datasets that can be easily converted to OME-Zarr or provide data in a format that is compatible with our on-demand transformer. More information on the on-demand transformer and our plan to serve OME-Zarr dynamically at BIL can be found in the publication describing community adoption of OME-Zarr^12^.

### Democratizing data analysis

Vast amounts of data generated by modern microscopes result in a data deluge, where analysis speed is far behind data generation speed. Furthermore, there are no agreed upon standard tools to analyze light microscopy brain images^36^, leaving many research groups to develop their own methods and image formats tailored to their data. This is often an unfortunate effort duplication, and many such tools are so esoteric that they never become reusable. The bioinformatics community may be a good model for how curated sets of analysis pipelines could be made available to the community^43^.

Considering that BIL is a centralized resource for the storage, sharing, analysis, and reuse of large brain imaging, it presents the optimal stage to further unify and consolidate whole brain analysis methods. By utilizing the data, the organizational structure and computational power offered at BIL, future projects can develop standardized and efficient analysis tools that benefit the neuroscience community at large. One use case for such tools may be to unify the thousands of whole brain datasets that exist at BIL into a simplified platform that enables high-level spatial meta analyses that crosses individual animals, experiments, laboratories and species. This in fact is a glaringly unmet need within the community. Whole brain data at BIL potentially contains billions of data points that describe cell positions, expression levels, cell morphologies and connectivity information. Some of this information exists latent within the BIL public archive, however, the majority of these higher-level features have never been extracted or have never been deposited. There needs to be a directed effort by the community to extract these features and to produce an online interface where a multitude of neuroscientists can ask novel questions, generate hypotheses and validate their results at the bench. We invite the neuroscience community to make use of BIL for exactly these types of efforts.

### Advancing FAIR image data practices

While public data resources focused on molecular biosciences have existed for over fifty years^44^, the culture of collecting and sharing image data is still in its early stages. Data has often resided at individual laboratories and has not been made public, or shared upon request. The change started with introduction of the FAIR principles in 2016^13^, which emphasized the importance of making all research data publicly available. The BRAIN Initiative’s data-sharing policy adopted these standards in 2019, which required all data being produced with BRAIN Initiative funding to be deposited in one of several designated archives^45^. In 2023, the NIH adopted a similar institution-wide data sharing policy^46^.

In line with the FAIR principles, BIL works towards improving data usability in multiple ways. Single-click visualization, accessible data analysis and serving multitude of image data formats are described above. However, as research in the life sciences embraces new technologies and becomes increasingly collaborative, it is important to establish linkages between resources such as ontological definitions, relevant anatomical atlases, and related data housed in different repositories. The potential value of the data stored at BIL, especially if cross-referenced and integrated across projects, is enormous. The first examples of this type of multi-modal, cross-scale imaging data integration already exist^47^. While this effort focused on mapping cell types in a mouse brain, the more recent BICAN consortium is focusing on mapping brain cell types in primates, including humans, presenting the possibility of cross-species comparisons. Therefore, our future efforts will focus on linking BIL data with other BRAIN Initiative resources and archives along with anatomical atlases produced by the community.

Connections between related data are facilitated by comprehensive metadata that references common searchable ontologies. Complicating this harmonization is that (i) each resource has evolved metadata schemas independently that serve their specific charters, thus there is no agreed-upon metadata standard that describes one-to-one relationships between resources, and (ii) annotation is an evolving target as cell types and atlases are refined and more fundamental knowledge is acquired. Where possible, we plan to systematically enhance our metadata with both automated and user-contributed annotations which can be easily searched and retrieved. As new imaging modalities are developed and widely adopted, BIL will continue to work with its community of collaborators to expand and refine the metadata to align with new technologies. Notably, BIL is currently engaged in a collaborative effort with BICAN members and subject matter experts to develop expanded metadata standards for spatial transcriptomics datasets. These datasets are expected to constitute a substantial portion of data deposits to BIL in the next several years.

We expect that beyond its primary objectives, BIL data can serve multiple purposes, including being used for developing novel analysis tools, benchmarking algorithms, improving visualization tools, teaching, and citizen science projects. Ultimately, we hope that ongoing work in the directions discussed above will facilitate collaboration, cross-archive integration, data access, and reuse, enhancing the value of BIL for data contributors and users.

## Data availability

All public data is available at BIL and searchable at https://brainimagelibrary.org, RRID: SCR_017272. DOIs and web links to publicly available datasets discussed herein have been included within the text.

## Code availability

All software products for BIL are developed in the open and stored in GitHub repositories https://github.com/brain-image-library and https://github.com/CBI-PITT. Jupyter notebooks for analyzing BIL data presented at workshops are available at https://github.com/brain-image-library/workshops. Source code for napari-bil-data-viewer is at https://github.com/brain-image-library/napari-bil-data-viewer. Code necessary to reproduce specific figures or visualizations are linked in the relevant figure legends.

## Acknowledgments

The Brain Image Library is supported by the National Institutes of Mental Health under award number R24-MH-114793. The napari visualization plugin is funded by the Chan Zuckerberg Initiative Donor Advised Fund grant #DAF2022-309651. This work utilized the Extreme Science and Engineering Discovery Environment (XSEDE), supported by National Science Foundation (NSF) award ACI-1548562, and allocation BIO210066 from the Advanced Cyberinfrastructure Coordination Ecosystem: Services & Support (ACCESS) program, supported by NSF awards #2138259, #2138286, #2138307, #2137603, and #2138296. Specifically, it used the Bridges system, supported by NSF award ACI-1445606, and the Bridges-2 system, supported by NSF award ACI-1928147. This content is solely the responsibility of the authors and does not necessarily represent the official views of the funding agencies mentioned.

The authors would like to thank the BIL development team (Kathy Benninger, Derek Simmel, Art Wetzel, Elizabeth Pantalone), Pittsburgh Supercomputing Centers Advanced Systems and Operations Group, the BIL Advisory Board (Jason Swedlow, Lydia Ng, Yongsoo Kim, Matt McCormick), Jesse F. Prentiss, Adam Tyson, BICCN and BICAN collaborators and our many data contributors.

## Author contributions statement

M.K. - writing, editing, reviewing, data curation; I.V. - writing, editing, reviewing, visualization, funding acquisition, G.H. - writing, editing, reviewing, I.C-B. - software, writing, editing, reviewing; L.T. - software, writing, editing, reviewing, R.L. - writing, editing, reviewing, M.S. - editing, reviewing, A.M.W. - conceptualization, supervision, writing, editing, reviewing, funding acquisition, visualization, methodology, A.J.R. - conceptualization, supervision, writing, editing, reviewing, funding acquisition.

## Competing interests

The authors declare no competing interests.

## Notes

### Competing Interest Statement

The authors have declared no competing interest.

### Summary of Updates

Add a results section about data analysis and a corresponding figure. Add a discussion section about democratizing data analysis. Merge several figures. Make the text more concise and readable. Rename several subsection headings.

